# Phenotypic plasticity is broadly adaptive across an elevation gradient in the Cutleaf Monkeyflower

**DOI:** 10.64898/2025.12.19.695179

**Authors:** Jill M. Love, Kathleen G. Ferris

## Abstract

- Phenotypic plasticity is a key mechanism by which organisms can cope with environmental heterogeneity, but its evolutionary consequences depend on how plastic responses align with the broader adaptive landscape.
- We tested whether plasticity in leaf shape alongside other traits is associated with differential fitness across an elevational gradient in *Mimulus laciniatus*, an annual wildflower endemic to montane California. Using a reciprocal transplant experiment and recombinant inbred lines (RILs) previously phenotyped for plasticity in controlled conditions, we measured variation in survival and fecundity in native low- and high-elevation habitats.
- Based on previous work, we expected to find selection against leaf shape plasticity at low-elevation and selection for leaf shape plasticity in the high-elevation direction (increased leaf lobing under long-day conditions) at high-elevation. Interestingly, we found that RIL genotypes exhibiting high-elevation plasticity had the greatest survival to seed production at high-elevations, but that low-elevation plasticity (increased leaf lobing under short-day conditions) was associated with greater fecundity in both elevations. RILs with high-elevation plasticity also outperformed non-plastic genotypes at high-elevation.
- This pattern suggests that plasticity is broadly beneficial across a species’ geographic range even if local variation in the direction of plasticity is not always adaptive.

## Introduction

In evolutionary ecology, many discussions have centered around how phenotypic plasticity affects an organism’s reproductive success across heterogeneous environments. Phenotypic plasticity, the ability of a single genotype to express multiple phenotypes depending on environmental conditions, offers organisms a degree of flexibility that may promote survival and reproduction when conditions shift (Bradshaw, 1965; West-Eberhard, 2003). By buffering against environmental variability, plasticity may enhance fitness in unpredictable or novel habitats, making it a potentially adaptive trait in the face of climate change or spatial heterogeneity (Via *et al*., 1995; Ghalambor *et al*., 2007). However, plasticity is not universally beneficial. It can incur physiological or genetic costs, such as resource allocation trade-offs, slower development, or increased susceptibility to herbivory or pathogens due to mismatched trait expression (Dewitt *et al*., 1998; Auld *et al*., 2010). Additionally, plastic responses that are beneficial in one environment may prove maladaptive in another, particularly if environmental cues are unreliable or plasticity limits local specialization (Gomulkiewicz & Kirkpatrick, 1992). These context-dependent effects make it difficult to generalize about the fitness consequences of plasticity. Despite decades of theoretical attention, empirical data on how plasticity influences fitness in the wild remains sparse (though see Scheiner & Callahan, 1999; Donohue et al., 2000; Dechaine et al., 2007; Hamann et al., 2025). This is partly due to logistical challenges: testing plasticity’s fitness effects requires genotypic replication across contrasting environments, a task often constrained by the difficulty of producing and transplanting clonal individuals into natural settings. As a result, many foundational questions remain unanswered, particularly regarding how plasticity is shaped by selection in complex and dynamic field environments.

Understanding plasticity’s role in local adaptation is especially interesting in environments that present steep ecological gradients. Altitudinal gradients in particular provide powerful natural laboratories to test for selection on phenotypic traits across contrasting environments because climate, abiotic stressors, and biotic interactions shift dramatically from low- to high-elevation within relatively short geographic distances (Körner, 2007; Halbritter *et al*., 2018). Across elevation, changes in temperature, growing season length, photoperiod, and moisture availability can create divergent selection pressures, driving population-level trait divergence. Plants are powerful models for studying adaptive plasticity because they are sessile organisms that must cope with environmental heterogeneity throughout their life cycle, often through developmentally flexible traits (Bradshaw, 1965; Sultan, 2000; Nicotra *et al*., 2010). Annual plants are particularly useful for evolutionary studies of plasticity because they experience only one reproductive season, allowing fitness to be measured comprehensively across environments in a single year (Schlichting & Smith, 2002; Donohue, 2003). In addition, their short generation times enable selection experiments across multiple environmental gradients. *Mimulus laciniatus,* the Cutleaf Monkeyflower, is a valuable model species for plasticity research, as it is a self-fertilizing, short-lived annual plant endemic to the Sierra Nevada (California, USA), growing across an elevational range from approximately 1,000 to 3,000 meters. In montane regions like the Sierra Nevada, lighter snowpack and earlier spring warming in low-elevation populations means these plants experience shorter days and higher temperatures, while high-elevation populations are characterized by cooler temperatures, higher snowpack, and shorter growing seasons under longer late spring/early summer photoperiods. These stark contrasts create divergent selection pressures that may favor different values of plasticity depending on elevation. In this study, we performed a reciprocal transplant experiment between low and high-elevation environments to test for selection on replicate recombinant genotypes varying in plasticity. This approach allows us to disentangle local adaptation from plastic responses by measuring trait-fitness associations in both native and non-native altitudes (Clausen *et al*., 1941; Ågren & Schemske, 2012).

In the genus *Mimulus*, trait divergence across environmental gradients is well documented, with evidence of selection on phenology, growth, and stress tolerance (Lowry *et al*., 2008; Hall *et al*., 2010; Sheth & Angert, 2016; Ferris & Willis, 2018; Kooyers *et al*., 2019; Tataru *et al*., 2023). Recent work in *M. laciniatus* demonstrated that populations differ not only in mean trait values across an elevational gradient, but also in the degree of leaf shape plasticity (Love & Ferris, 2024). In *M. laciniatus*, high-elevation populations exhibit greater leaf lobing in response to increased photoperiod, whereas low-elevation populations exhibit little leaf shape plasticity, or plasticity in the opposite direction where leaves become more lobed under shorter photoperiods. This difference corresponds to elevation-specific environmental cues, specifically the timing of snowmelt causing high-elevation populations to germinate and develop later under longer photoperiods closer to the summer solstice than low-elevation *M. laciniatus*. These findings suggest that plasticity itself may respond to locally varying selection, potentially contributing to local adaptation. While Love & Ferris (2024) provided strong evidence for divergent plastic responses in controlled environments, whether those plasticity differences influence fitness in the wild remains untested. The present study builds on previous work, using a newly created recombinant inbred line (RIL) population and field-based approach to ask whether plasticity in *M. laciniatus* contributes to local adaptation across elevations.

By measuring multiple phenotypic traits, we are able to capture a fuller picture of how selection acts across different functional axes of plant growth. Given well established links between plant size, height, flowering time, and fitness in annual species, we predict that RILs exhibiting larger size, taller growth, and earlier flowering will have higher fitness (Mitchell-Olds, 1996; Stanton *et al*., 2000; Hall & Willis, 2006; Franks *et al*., 2007; Wadgymar *et al*., 2017). We also predict that the direction and magnitude of selection on plastic genotypes will vary based on environmental context, as demonstrated in field studies across plant taxa. Studies have shown that phenotypic plasticity can be under environment dependent selection and can confer benefits or costs depending on context. For example, in *Impatiens capensis*, selection favored plastic shade-avoidance traits in dense understory sites, but not in open, low-density habitats (Donohue *et al*., 2000), demonstrating that the fitness benefits of plasticity may emerge only under particular environmental constraints. In *Brassica rapa*, plasticity incurred detectable fitness costs under high-density conditions, further illustrating that plastic responses are not universally beneficial (Dechaine *et al*., 2007). In *M. laciniatus*, leaf shape is an interesting trait to examine in the context of plasticity and local adaptation, as it is known to influence key plant physiological processes. More highly lobed leaves reduce boundary layer thickness, improving convective heat loss in hot environments (Gurevitch & Fox, 2006; Nicotra *et al*., 2011). Lobed leaf morphology may also enhance hydraulic efficiency and reduce resistance to water transport, allowing improved transpiration in dry or stressful environments (Nicotra *et al*., 2011). These functional tradeoffs suggest that selection on leaf shape plasticity may depend on specific challenges posed by each environment, and that locally adaptive plasticity in leaf lobing could be a key component of *M. laciniatus*’ broad elevational distribution.

To test for locally varying selection on phenotypic plasticity, we generated a recombinant inbred line (RIL) population from low- and high-elevation *M. laciniatus* genotypes exhibiting different degrees of leaf shape plasticity. This RIL population allowed us to ask whether plasticity is broadly or locally adaptive across an elevational gradient of *M. laciniatus* native habitat. If plasticity is broadly beneficial in buffering against environmental variation in this species, we expect positive selection for more plastic RIL genotypes across both high and low-elevation sites. However, if plasticity imposes costs or only confers advantages under specific conditions (i.e. is locally adaptive with trade-offs), the direction and magnitude of selection on plastic genotypes will vary depending on the environmental context. Based on previously observed plasticity clines in Love and Ferris (2024), we expect to find selection against plasticity in low-elevation environments and selection for high-elevation plasticity (increased leaf lobing in long days) at high-elevations. By measuring genotypic selection on 4,950 individuals across a reciprocal transplant with five common garden field sites, we tested whether plasticity in leaf shape, along with mean variation in leaf shape, height, and flowering time, is locally adaptive across *M. laciniatus’s* altitudinal range.

## Materials and Methods

## Creation of Recombinant Inbred Lines

To measure phenotypic selection on a genetically replicable population, we generated RILs that segregate in leaf shape plasticity expression. We created a population of 225 *M. laciniatus* RILs by crossing a low-elevation, non-plastic *M. laciniatus* genotype (HEH 20, Huntington Lake, CA) with a high-elevation, highly plastic *M. laciniatus* genotype (WLF 72, White Wolf Campground, CA) to make an F_1_ hybrid (Fig. **1a**). We then self-fertilized a single F_1_ hybrid to create 500 F_2_ individuals, self-fertilizing each line to the F_8_ generation (Pollard, 2012). Over the course of RIL creation we lost 55% of lines to inbreeding depression. A crossing diagram depicting the details of RIL design can be found in Fig. S1. Due to logistical constraints and rapid drying of our field transplant sites, we were unable to consistently obtain high-quality leaf shape measurements across all genotypes and elevations in field conditions without damaging the plants and affecting fitness. As a result, we used phenotypic data from a common garden growth chamber experiment to represent genotypic values of trait plasticity, and data from a separate common garden greenhouse experiment to quantify genotypic mean traits in each RIL. These common garden experiments are described in detail in the next paragraph. This approach aligns with established genotypic selection frameworks that estimate selection on traits using average trait values per genotype (Rausher, 1992; Stinchcombe & Rausher, 2001). Due to logistical constraints mentioned above, our approach differs slightly in that trait means were measured in a common environment while fitness was measured in the field. This design enables us to evaluate whether trait plasticity, as an inherent property of a genotype, correlates with fitness in natural environments.

**Figure 1.**
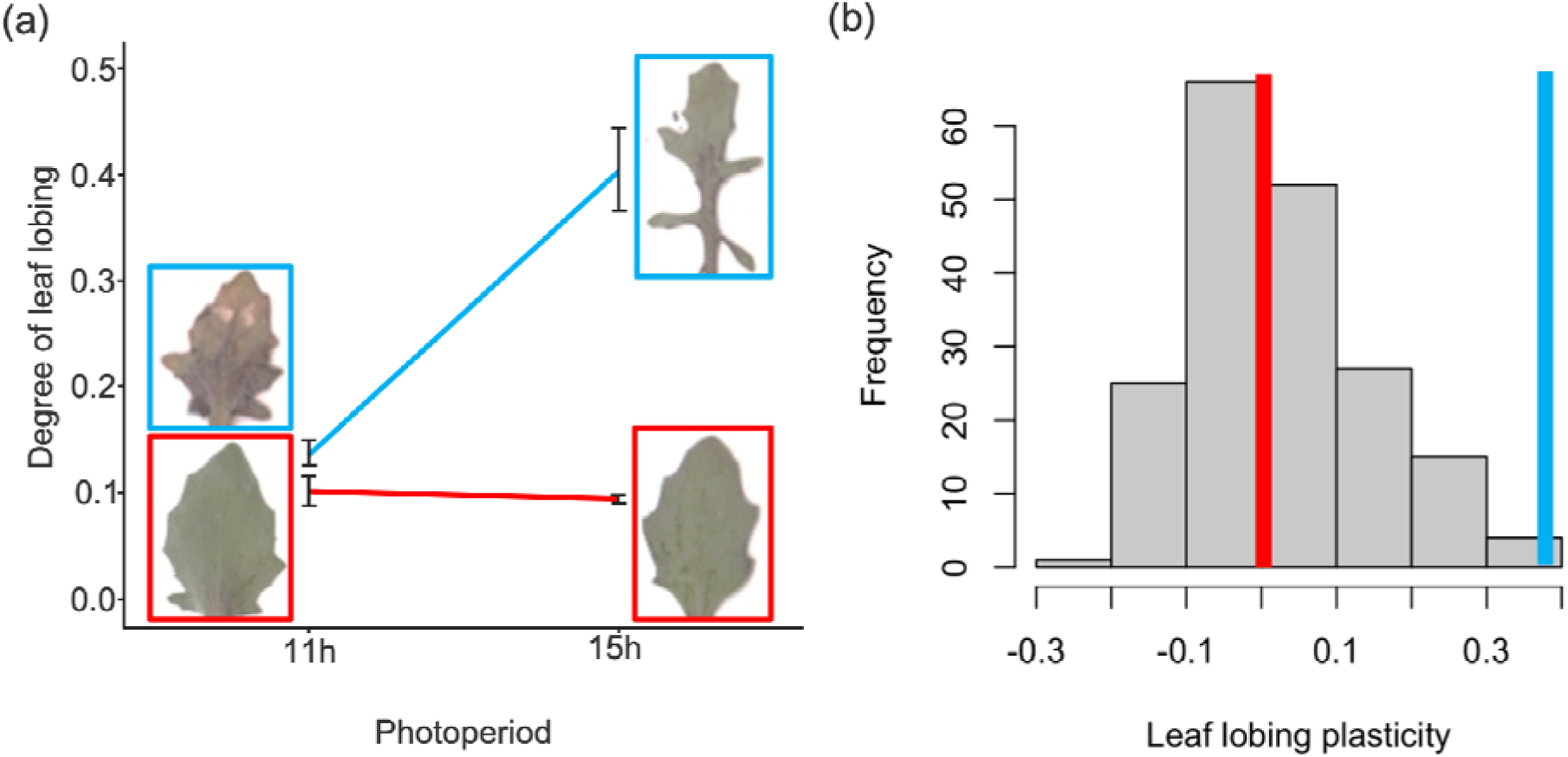
**a**) Leaf shape reaction norms of low- and high-elevation *M. laciniatus* parental genotypes used in RIL creation in low (11h) and high-elevation (15h) photoperiods. Error bars represent standard error. **b**) Distribution of leaf lobing plasticity expression in the RIL population (gray bins) relative to genotypic plasticity values of the low- (red) and high-elevation (blue) parental lines used in RIL creation.

### Measuring genotypic values of mean and plastic RIL traits

To measure the genotypic value of plasticity expressed by each genetic line, we grew three replicates of each genotype and both parental genotypes under 11- and 15-hour photoperiods in a Conviron growth chamber and compared them to mean leaf shape values measured in a separate greenhouse experiment as described in Love & Ferris, 2024. This design avoids the statistical artifact of regression to the mean (Gunderson & Revell, 2022). On the day of first flower, we recorded the date to calculate flowering time, measured plant height, and collected the first true leaf from all plants for leaf lobing analysis in ImageJ (Ferris *et al*., 2015). Briefly, the lobing index provided by leaf lobing analysis is a metric from zero to one, calculated by dividing the difference between the true area of the leaf and the larger area of the leaf’s convex hull by the area of the convex hull. Plasticity per genotype was calculated as the mean 15-hour phenotype subtracted from the mean 11-hour phenotype; the full distribution of RIL leaf shape plasticity is displayed in Fig. **1b**. The expression of leaf shape plasticity in the RILs is transgressive relative to the low-elevation parent, where we found significantly more negative values of plasticity than our low-elevation parent, which had near-zero plasticity. In our previous common garden experiment of *M. laciniatus* from across the species’ range, many low-elevation genotypes did have negative values of plasticity, meaning these genotypes developed more lobed leaves under short days (Love and Ferris 2024). Therefore, we consider RILs with a negative value of plasticity to have low-elevation-like plasticity and will use that terminology throughout. To quantify the genotypic mean expression of flowering time, plant height, and leaf lobing, we again grew three replicates of each RIL genotype and both parental genotypes under 16-hour photoperiod in the Tulane University greenhouse, phenotyping as described above.

### Field reciprocal transplant experiment

At the outset of our field reciprocal transplant experiment in 2024, RIL seeds were planted in 78-cell trays with Fafard 4B soil, kept moist and dark in stratification at 4°C for 10 days. Trays were then moved to the UC Merced greenhouse and misted daily until germination. Plants stayed in the greenhouse for 7-10 days before transporting to field sites for planting. Greenhouse germination occurred twice during the experiment, for low-elevation sites on March 24 and high-elevation sites on May 28. We chose five common garden locations in native *M. laciniatus* habitat within Yosemite National Park, CA – three at low-elevation (LMG, TRT, TNL) and two at high-elevation (YCN, OPN) (Table 1). This work was permitted by the National Park Service. Planting began when snow melted and native *M. laciniatus* began to germinate. To plant each reciprocal transplant site, all native angiosperm plants were first cleared, then we transplanted four replicates of each RIL genotype and 100 replicates of each RIL parental genotype at the cotyledon stage into 44 blocks of a randomized block design (25 individuals per block in 5x5 array) for a total of 1100 plants per site. One low-elevation site, TNL, had half as many blocks and replicates. One week after planting was completed, dead transplants were re-planted to account for transplant shock. Low-elevation planting finished on April 11 (except TNL, which due to strong snowmelt runoff finished on June 4) and high-elevation on June 11. Afterward, sites were monitored every three days for survival and flowering. On the date of first flower, plant height was measured. After plants senesced, we collected all fruits and counted the total seed number for each plant as our fitness metric.

**Table 1.** Geographic information for each reciprocal transplant site. Sites were grouped into low or high-elevation categories, with exact elevations and GPS locations provided.

| Site | Elevation (meters) |  | GPS coordinates |
| --- | --- | --- | --- |
| Tunnel View (TNL) | Low | 1347.0 | 37.71482, -119.67715 |
| Turtleback Dome (TRT) | Low | 1485.2 | 37.71594, -119.70528 |
| Lower Mariposa Grove (LMG) | Low | 1625.1 | 37.49959, -119.62682 |
| Yosemite Creek (YCN) | High | 2325.6 | 37.84369, -119.57309 |
| Olmsted Point (OPN) | High | 2563.6 | 37.80720, -119.48636 |

### Statistical Analyses

All statistical analyses were performed in R v4.5.1 (R Core Team, 2025). We calculated average genotypic fecundity as the total seed number divided by the total number of plants planted for each genotype in each habitat. To detect patterns of local adaptation in parental genotypes, we also calculated survival, as the proportion of individuals from each parental genotype that survived to flowering at each transplant elevation from the number of replicates planted. Additionally, we calculated parental total fitness as the number of seeds per planted replicate. Correlation matrices of low- and high-elevation traits were created using the *corrplot* package in R (Wei & Simko, 2021).

#### Environmental variation

In order to quantify how survival varied across elevations based on environmental context, we collected weekly fine-scale environmental data on soil moisture (Dynamax SM150 Soil Moisture Sensor) and soil surface temperature (laser thermometer). We then created linear mixed-effects models using the *nlme* package in R (Pinheiro *et al*., 2023). Survival was calculated weekly as the proportion of individuals alive within each block, and modeled as a function of time, site, soil moisture, and surface temperature (as in Ferris & Willis, 2018; Tataru *et al*., 2023; Dong *et al*., 2025). Dependent variables were treated as fixed effects, and block was included as a random effect. Our model was fitted using maximum likelihood (ML) to enable comparison among models with different fixed-effect structures. We used the *dredge* function in the *Mu-MIn* package (Barton, 2009) to perform model selection. *Dredge* evaluates all possible subsets of the global model and ranks them by Akaike Information Criterion (AIC). We extracted the best-supported model based on highest AIC. Finally, to assess the contribution of each predictor in the model, we conducted a marginal ANOVA using Type III sums of squares.

#### Local adaption and genotypic selection analysis

To understand the relationship between measured plant traits and fecundity we performed genotypic selection analysis using general linear models in the R package *GlmmTMB* (Brooks *et al*., 2017; R Core Team, 2025). We analyzed plant fecundity in parental genotypes as a means to test for local adaptation at low and high-elevation common gardens. To analyze RIL data, we included field-measured traits as well as genotypic means and genotypic plasticity expression from lab-collected phenotypic measurements as independent variables in our models, and field-measured averaged seed number per genotype (genotypic fitness) as our dependent variable, with block nested in site as a random effect. Our full list of traits as independent variables encompassed plant height (field and genotypic mean), flowering time (field and genotypic mean), leaf lobing (genotypic plasticity and mean), and leaf area (genotypic plasticity and mean). Because of limited survival in each individual field site we combined datasets into two groups, low-elevation and high-elevation (Table 1), running full models with all traits in each elevation dataset separately. Trait values were standardized across datasets to allow direct comparison of selection gradients.

To deal with an excess of zeros in our fitness data we used a hurdle model approach (Anderson *et al*., 2015; Ferris & Willis, 2018; Tataru *et al*., 2023). We broke the data into two subsets: 1) all individuals that flowered and 2) all individuals that flowered and produced at least one seed. We used a multivariate binomial model to assess whether a trait is associated with survival to producing seed (data subset 1), with a fecundity of either 0 or 1 as the dependent variable. For data subset 2, we used a zero-truncated Poisson regression as a measure of whether each trait is associated with the number of seeds produced, with fecundity as the dependent variable. All models can be found in Table 3. For both data subsets, we performed additional model selection using *Mu-MIn* in R to determine the best-fit model in each elevation environment (Barton, 2009) which can be found in Table S1. To visualize results of the zero-truncated Poisson models, we used the R package *visreg* (Breheny & Burchett, 2017).

We were interested in whether phenotypic plasticity confers a broad or locally adaptive advantage. We ran an ANOVA on a linear model to test whether different directions of leaf shape plasticity were associated with distinct fitness outcomes across both altitudinal habitats. To do this, we combined the low- and high-elevation datasets into one and categorized plasticity expression into three groups: low-elevation, where expression equalled <–0.1; none, where expression equalled ±0.1 from zero; and high-elevation, where expression equalled >0.1. Fecundity was the dependent variable, with plasticity group as the independent variable and site as a random effect. Phenotypically, negative values of leaf shape plasticity (low-elevation plasticity) indicated that genotypes expressed more deeply lobed leaves under short days, which was the most common direction of expression in *M. laciniatus* genotypes from low-elevation environments in Love and Ferris (2024). On the other hand, positive values (high-elevation plasticity) occur when genotypes express more deeply lobed leaves under long-day photoperiods typically only experienced by high-elevation populations. To understand the direction of selection on plasticity within each low- and high-elevation habitat, we used the R package *emmeans* (Lenth, 2025) to conduct post-hoc asymptotic z-tests on our zero-truncated Poisson models to compare model-adjusted fitness across the same three categories of leaf shape plasticity.

## Results

### Time, site, soil moisture, and soil temperature all significantly influence survival

Survival declined strongly over the course of the season and was shaped by multiple environmental gradients (Fig. 2, Table 2). Days since planting was the strongest predictor of survival (F_1312_ = 406.27 p < 0.0001), indicating a sharp temporal decrease in the proportion of surviving plants (Fig. **2a**). This temporal decline was further modified by soil moisture, with a highly significant Time × Soil moisture interaction (F_1312_ = 358.42, p < 0.0001) and Time × Soil temperature interaction (F_1312_ = 23.27, p < 0.0001), showing that the effect of time on survival depended on moisture and temperature conditions. Standalone environmental variables also showed strong effects on survival. As found in previous field experiments in *M. laciniatus* habitat (Ferris and Willis 2018; Tataru et al. 2023; Dong et al. 2025), soil moisture (F_1312_ = 36.78, p < 0.0001; **Fig. 2bc**) and soil temperature (F_1312_ = 42.18, p < 0.0001; **Fig. 2cb**) each significantly influenced survival.

**Figure 2.**
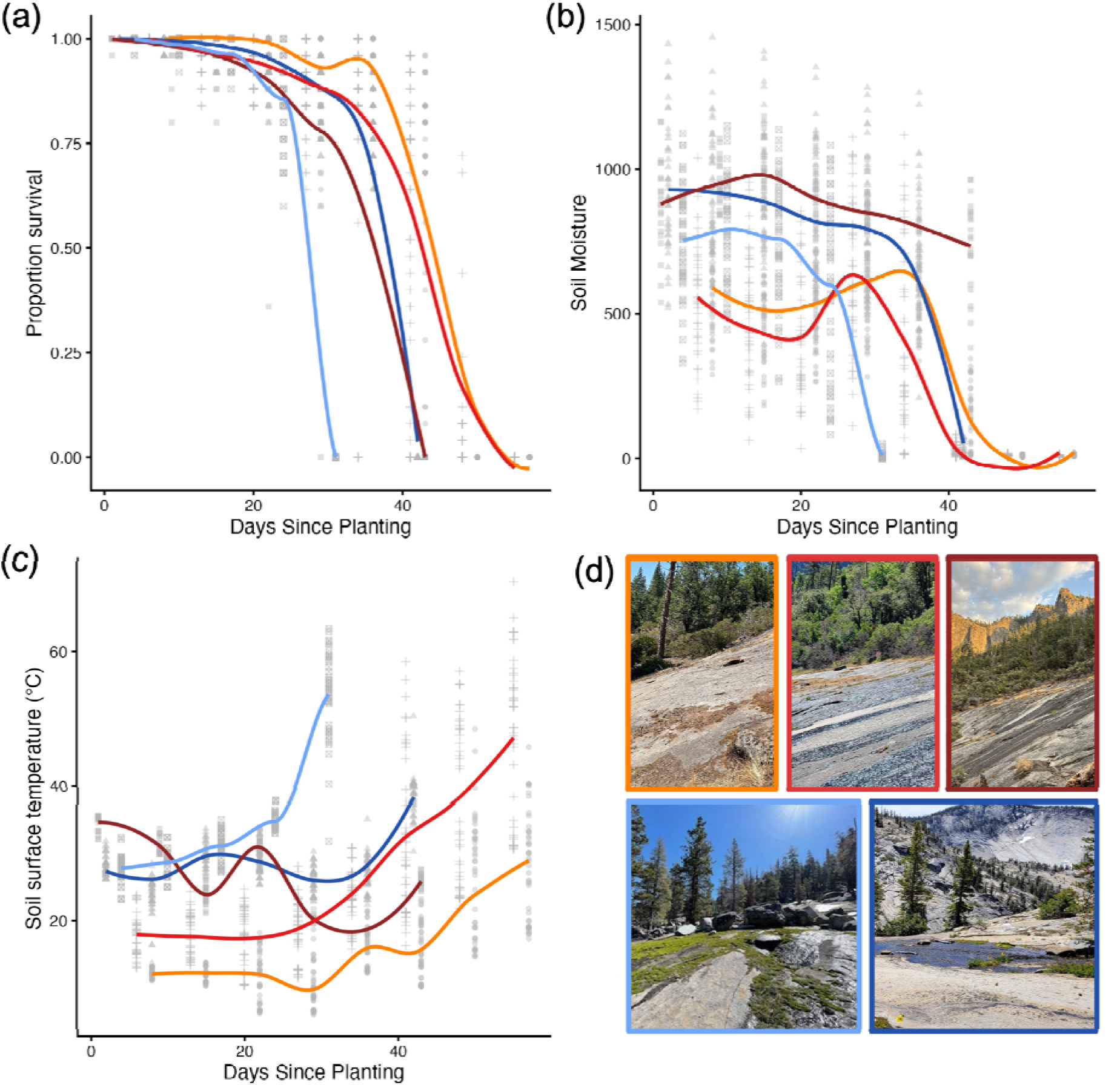
Survivorship curves for experimental plants in low-elevation (red and orange) and high-elevation (blue) common gardens. We display **a**) proportion survival, **b**) soil moisture, and **c**) soil surface temperature over time (Table 2). Panel **d**) shows a photo of each site.

**Table 2.**
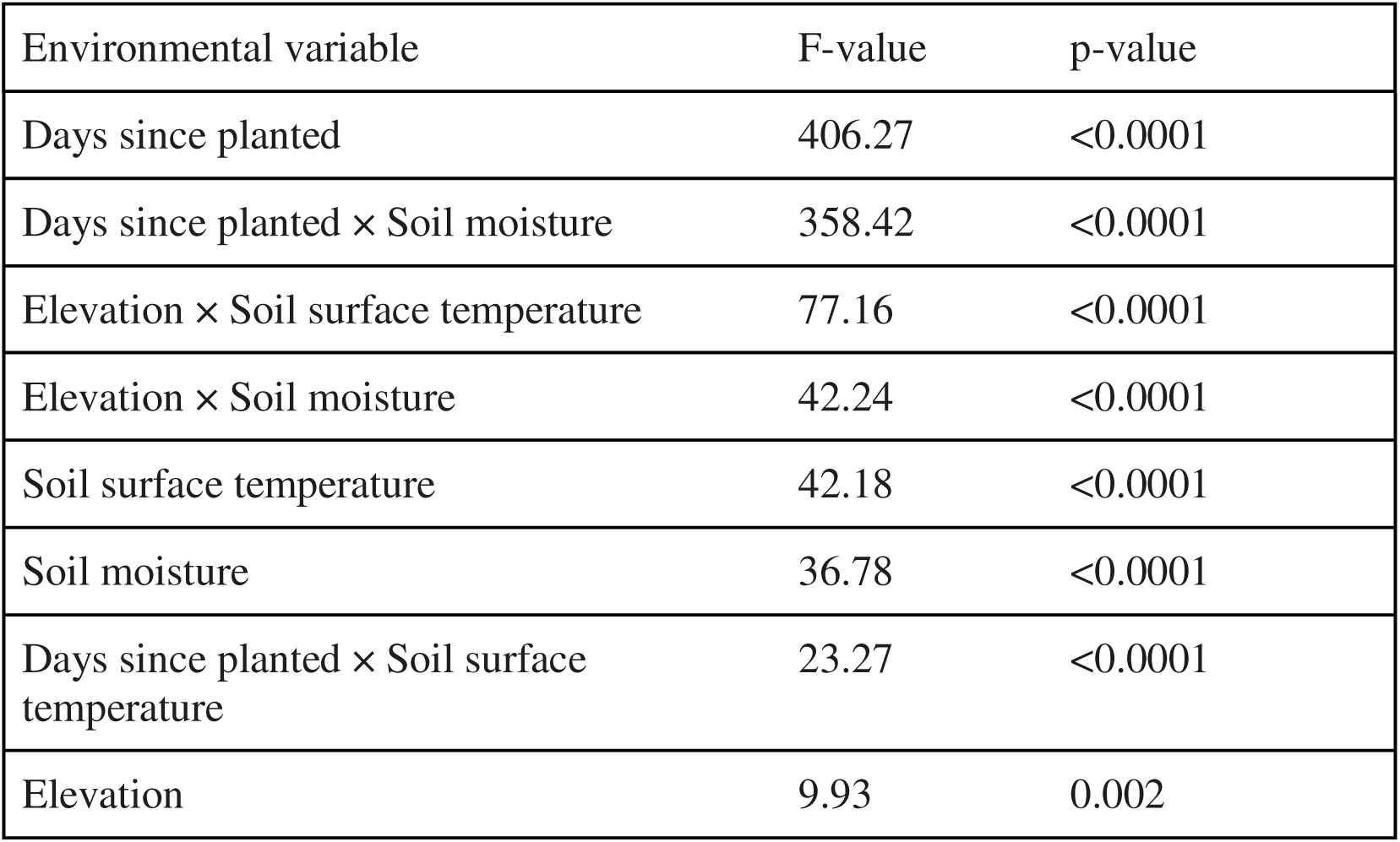
Results of ANOVA on a linear model investigating the impact of measured environmental variables on plant survival along the duration of the reciprocal transplant. For all environmental variables, numDF = 1, denDF = 1312.

| Environmental variable | F-value | p-value |
| --- | --- | --- |
| Days since planted | 406.27 | <0.0001 |
| Days since planted × Soil moisture | 358.42 | <0.0001 |
| Elevation × Soil surface temperature | 77.16 | <0.0001 |
| Elevation × Soil moisture | 42.24 | <0.0001 |
| Soil surface temperature | 42.18 | <0.0001 |
| Soil moisture | 36.78 | <0.0001 |
| Days since planted × Soil surface temperature | 23.27 | <0.0001 |
| Elevation | 9.93 | 0.002 |

**Table 3.**
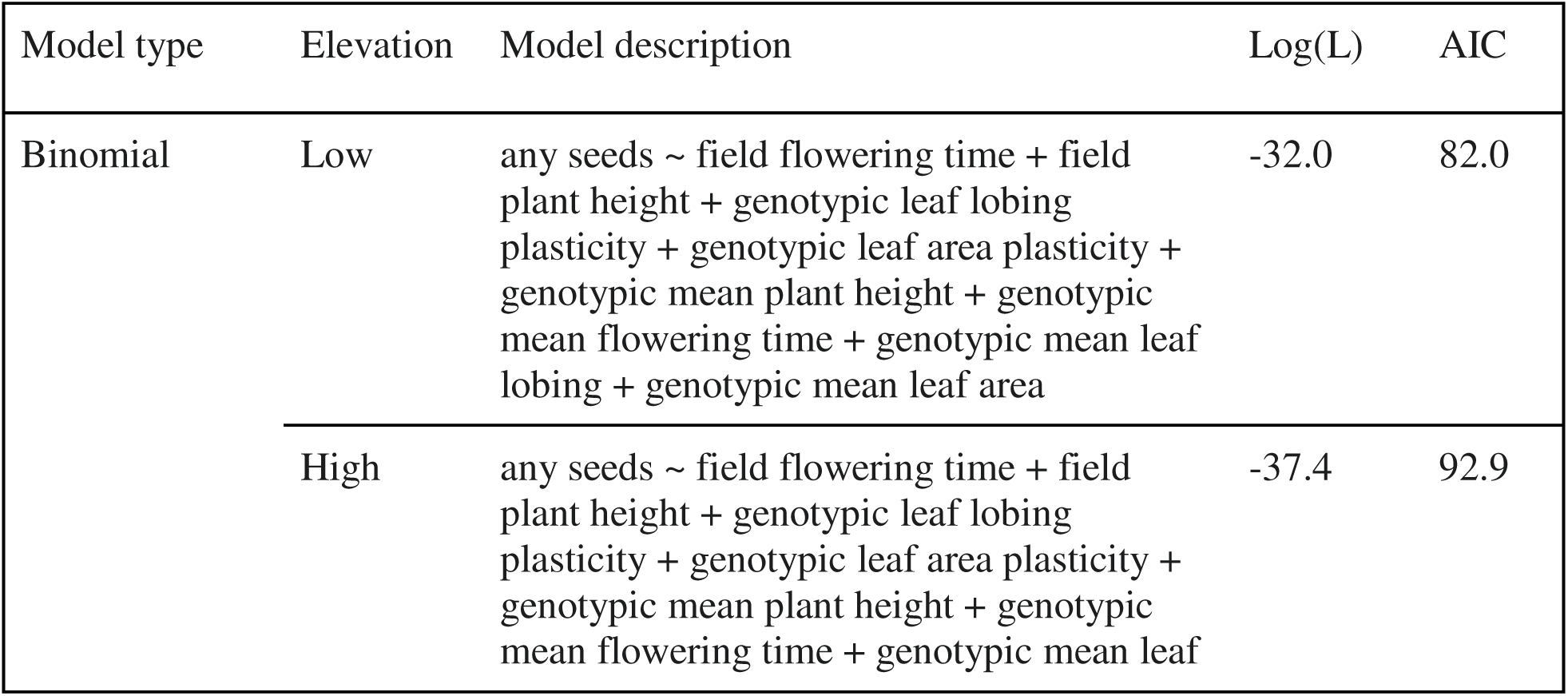
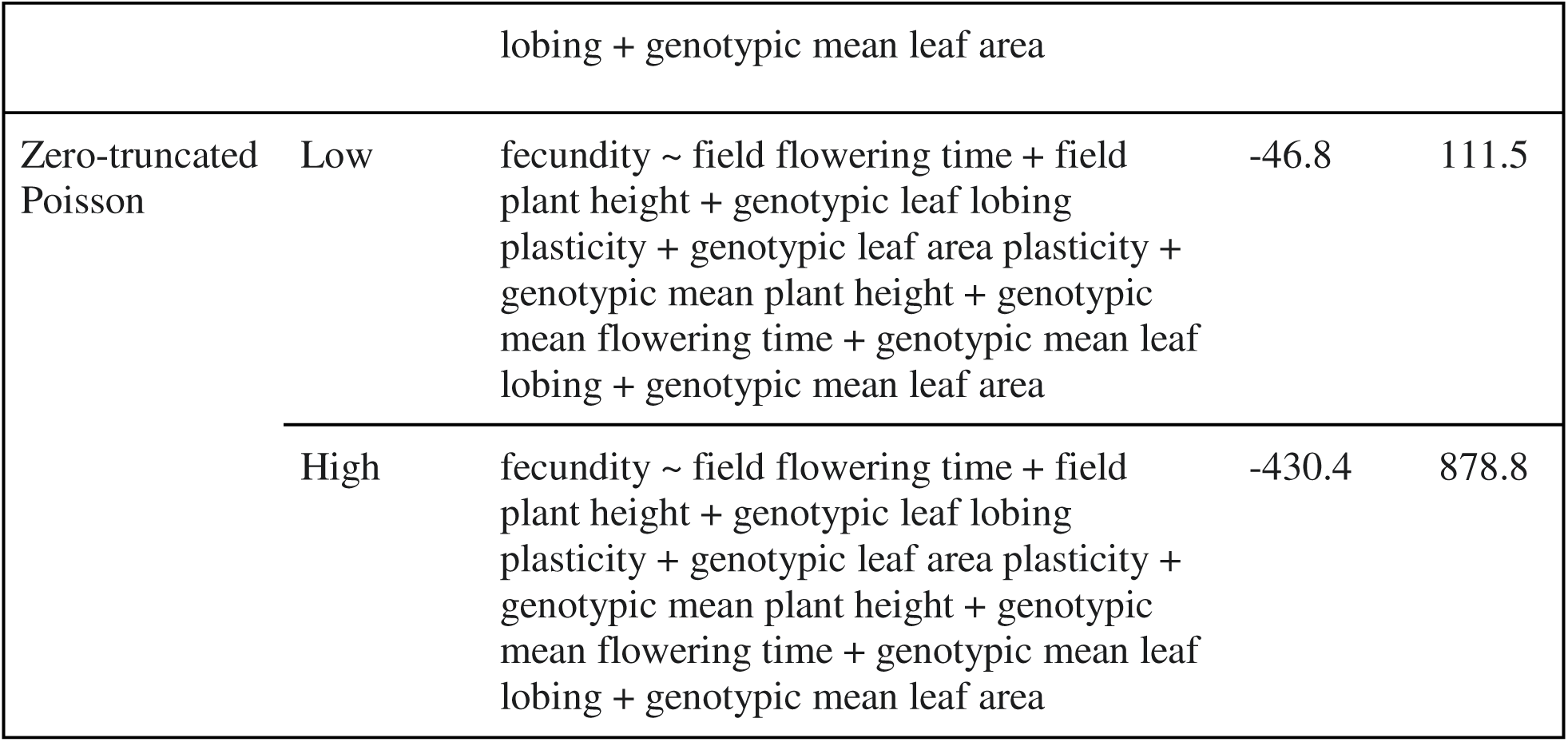
Comparison of binomial and zero-truncated Poisson models for low- and high-elevation sites. All predictor variables were included in each model, and log-likelihood (log(L)) and AIC are provided to assess relative model fit.

Survivorship curves varied significantly among elevations, with more prolonged survival at low-elevation compared to high (F_1312_ = 9.93, p = 0.002). We found that Elevation × Soil moisture had similar modulating effects on survival (F_1312_ = 42.24, p < 0.0001). As expected, soil moisture was higher at the beginning of the growing season at high-elevation sites than it was at two out of three low-elevation sites (Figure **2b**). The third low-elevation site, TNL, contained a continuous waterfall that did not dry up even late in the season. Elevation-level differences also shaped thermal and hydrologic effects. Elevation × Soil temperature had a significant impact on survival (F_1312_ = 77.16, p < 0.0001), demonstrating that the impact of temperature on survival varied across the environmental contexts represented at each elevation. Unexpectedly, we found higher soil temperature at the beginning of the season in high-elevation sites compared to two out of three low-elevation sites (Figure **2c**). This is likely because we had to delay planting at high-elevation sites until later in the season due to the timing of snow melt. However, we see that the seasonal increase in soil temperature was greater at low-elevation sites than high, falling in line with our expectations. Together, these results highlight that abiotic stressors such as heat and drought exert different selective pressures across elevations. Environmental context, particularly elevation-mediated variation in moisture and temperature, modulates both the pace and magnitude of mortality in this system.

### The low-elevation parent shows higher fitness across environments

Parental genotypes exhibited distinct reaction norms in survival, fecundity, and total fitness across transplant elevations, but these patterns did not align with classic expectations of local adaptation with trade-offs. Local adaptation is supported when each genotype has the highest fitness in its home environment relative to a foreign genotype (Kawecki & Ebert, 2004). To assess this, we compared the survival, fecundity, and total fitness of the low- and high-elevation parental genotypes within each elevation. We first assessed parental survival to flowering to determine whether genotypes exhibited differences in viability across elevations (Fig. **3a**). At low-elevation, both parental genotypes exhibited very low survival to flowering, with low parents having slightly, but not significantly, greater fitness in their local environment (low parent: 0.8%; high parent: 0%). In contrast, at high-elevation, the low-elevation parent had substantially greater survival (23.0%) than the high-elevation parent (3.5%), with an estimated odds ratio of 0.10 (95% CI: 0.03–0.25) indicating roughly a tenfold difference in survival. This difference was strongly supported by a Fisher’s exact test (*p* < 0.0001). These results show that contrary to predictions under local adaptation with trade-offs (Kawecki & Ebert, 2004), the low-elevation genotype outperformed the high-elevation genotype at both sites and did better at its non-native site when comparing within genotype across elevations.

**Figure 3.**
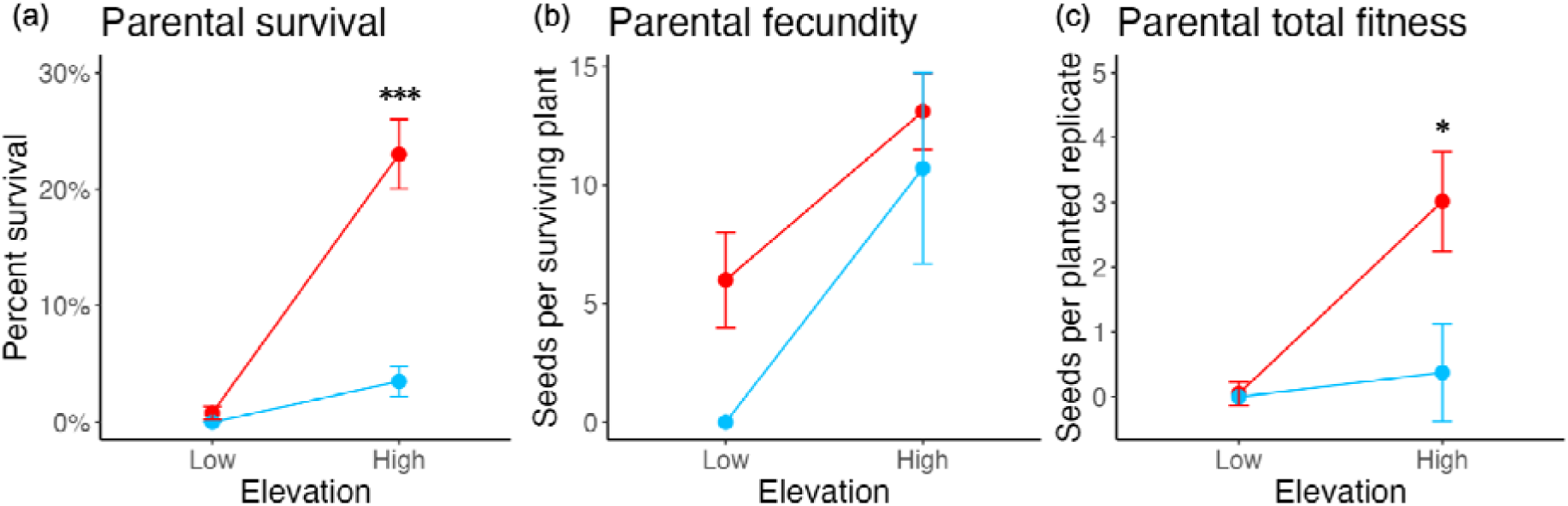
*Mimulus laciniatus* high (blue) and low-elevation (red) parental genotype reaction norms in **a**) survival, **b**) fecundity, and **c**) total fitness across experimental garden elevations. Error bars indicate standard error and statistical significance is indicated as * p-value < 0.10, ** p-value < 0.05, *** p-value < 0.01

Among plants that survived to reproduction, we examined fecundity per surviving individual, allowing us to separate survival effects from variation in seed production (Fig. **3b**). At low-elevation, the low-elevation parent produced 6.00 seeds per plant on average, but low sample size meant this was not significantly different from the high-elevation parent’s zero seed production (t_1_ = 3.0, p = 0.205). At high-elevation, both genotypes produced similar numbers of seeds with the low-elevation parent having slightly higher fecundity (low parent: 13.109 seeds; high parent: 10.714 seeds). This indicates that once plants survive to flower at high-elevation, fecundity between genotypes does not differ significantly (t_8_ = 0.552 *p* = 0.596).

Finally, we combined survival and fecundity to estimate total parental fitness as the number of seeds per planted individual, providing an integrated measure of lifetime reproductive success across elevations (Fig. **3c**). At low-elevation, both genotypes had near zero total fitness, but with the low parent again having a slightly higher value (low parent: 0.048 seeds, high parent: 0 seeds; Wilcoxon rank-sum test: W = 31500, *p* = 0.158). At high-elevation, the low-elevation parent produced 3.015 seeds per planted replicate, while the high-elevation parent produced 0.375 seeds (Wilcoxon rank-sum test: W = 23934, *p* < 0.0001). Overall, these results suggest that the low-elevation genotype consistently outperformed the high-elevation genotype across elevations.

### Both mean and plastic traits are under genotypic selection at high and low-elevations

To evaluate which traits most strongly predicted survival to flowering across environments, we used a genotypic selection analysis with multivariate binomial models for low- and high-elevations (Table 4). These models included combinations of field-measured traits, genotypic mean traits, and genotypic values of plasticity to evaluate their relative contributions to fecundity. Best-fit models may be found in Table S1, and a phenotypic correlation matrix can be found in Fig. S2. At low-elevation, only field-measured plant height was under marginally significant positive selection in the binomial model (β = +0.706, z_171_ = 1.781 p = 0.075). At high-elevation, more traits in the binomial model were under significant directional selection: taller field plant height (β = +1.903, z_169_ = 3.077, *p* = 0.002), earlier field flowering time (β = -0.761, z_169_ = -2.213, p = 0.027), and high-elevation (positive slope) genotypic leaf lobing plasticity (β = +0.762, z_169_ = 1.835, p = 0.067) all led to a higher likelihood of surviving to seed production. These results suggest that viability selection at high-elevation favors taller genotypes with earlier flowering and increased plasticity in the local high-elevation direction, while the sole trait predictor for survival to seed production at low-elevation, plant height, was under weaker and marginally significant selection.

**Table 4.**
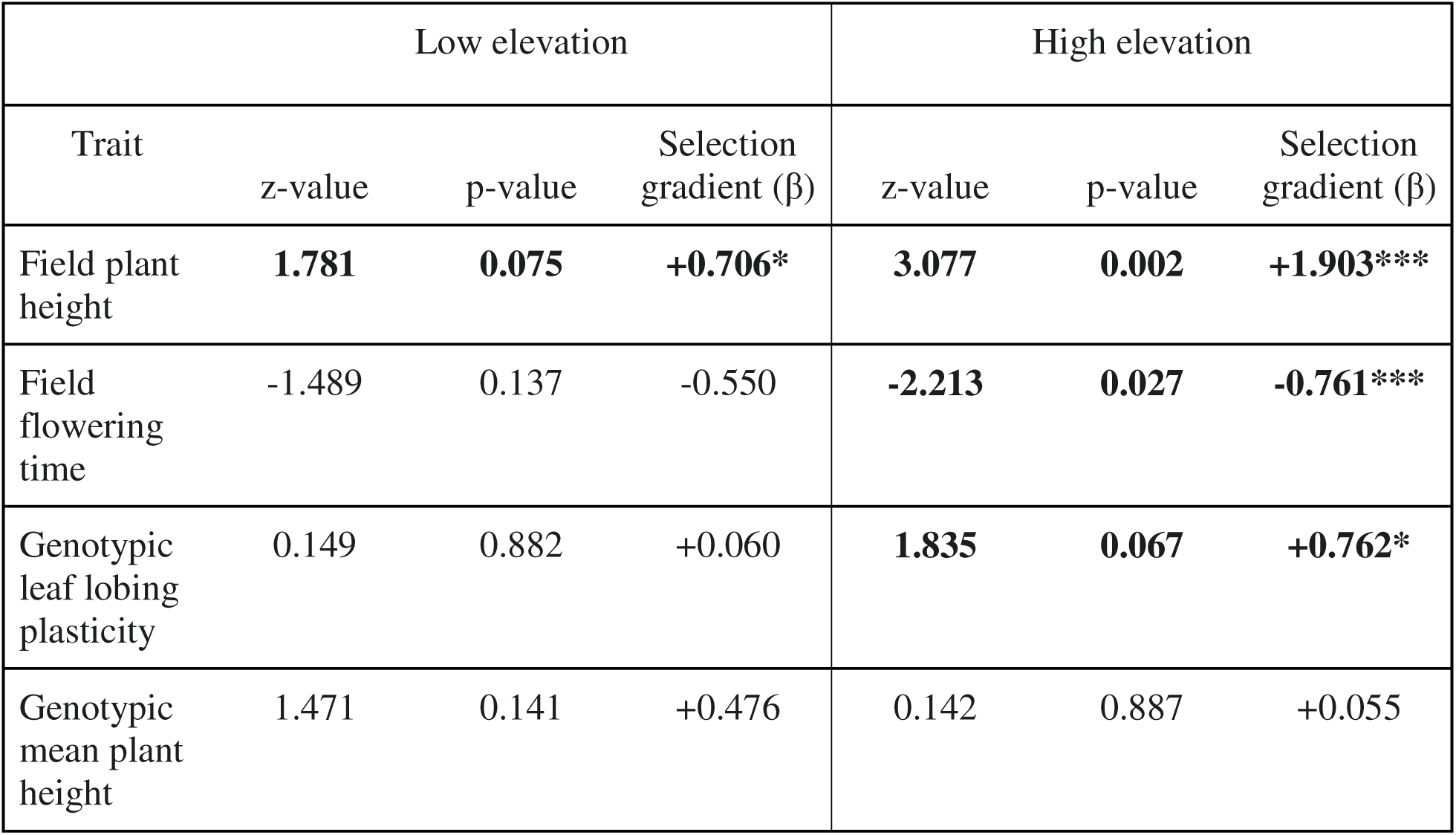
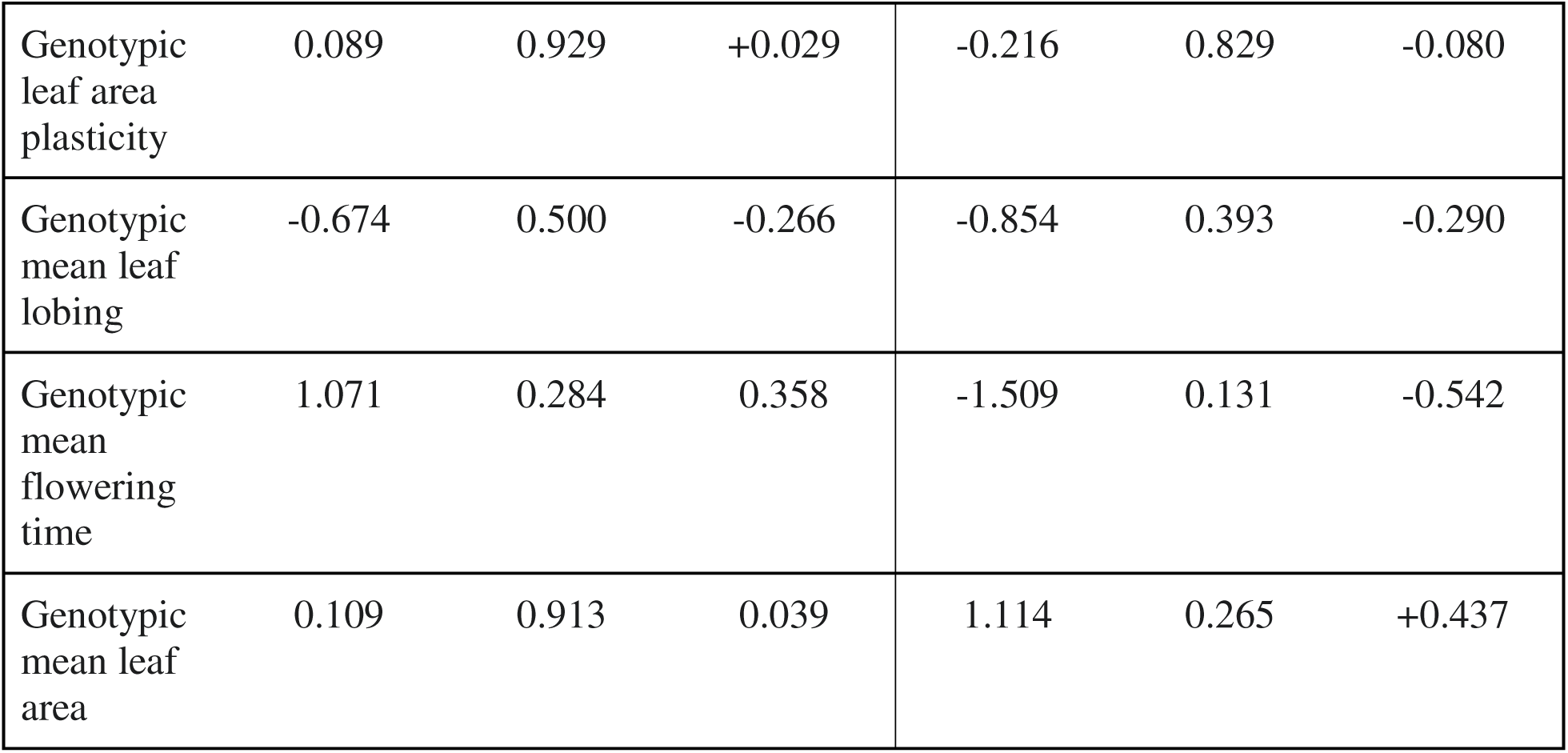
Trait influence on presence of seed production, based on binomial models run on low-and high-elevation datasets. Selection gradient (β) represents the strength and direction of selection on a trait. For low-elevation traits, numDF = 1, denDF = 171. At high-elevation, numDF = 1, denDF = 169. * p < 0.10, ** p < 0.05, *** p < 0.01

To identify specific traits under fecundity selection across environments, we fit zero-truncated Poisson models using genotype-level data at each site. At low-elevation, significant variables in the model included field-measured plant height, genotypic mean plant height, and genotypic leaf shape plasticity (Table 5, Fig. 4). Taller plants, measured both as genotypic mean and in the field, had significantly positive selection (field height: β = +0.690, z_18_ = 2.556, p = 0.011, genotypic mean height: β = +0.302, z_18_ = 1.968, p = 0.049). Leaf shape plasticity was under significant directional selection that favored local, low-elevation plasticity expression (β = -0.451, z_18_ = -2.432, p = 0.015). Interestingly, we also found significant positive selection for mean leaf lobing (β = +0.326, z_18_ = -2.101, p = 0.036). At high-elevation, fecundity was also predicted by a broad suite of traits (Table 5, Fig. 4). Field-measured flowering time was under negative selection, indicating selection for an earlier flowering time (β = -0.282, z_96_ = -5.75, p < 0.001), while taller field plants (β = +0.289, z_96_ = 7.07, p < 0.001), greater genotypic mean leaf lobing (β = +0.088, z_96_ = 2.58, p = 0.010), and greater plasticity in leaf area were all under positive selection (β = +0.115, z_96_ = 3.11, p = 0.002). Leaf lobing plasticity was also under marginally significant directional selection favoring the low-elevation phenotype (β = -0.070, z_96_ = -1.68, p = 0.093). This result was somewhat surprising given that low-elevation plasticity, producing more lobed leaves under short-day conditions that mimic the photoperiod of low-elevation sites, was not the previously observed dominant phenotype in local genotypes at high-elevation sites (Love & Ferris, 2024).

**Figure 4.**
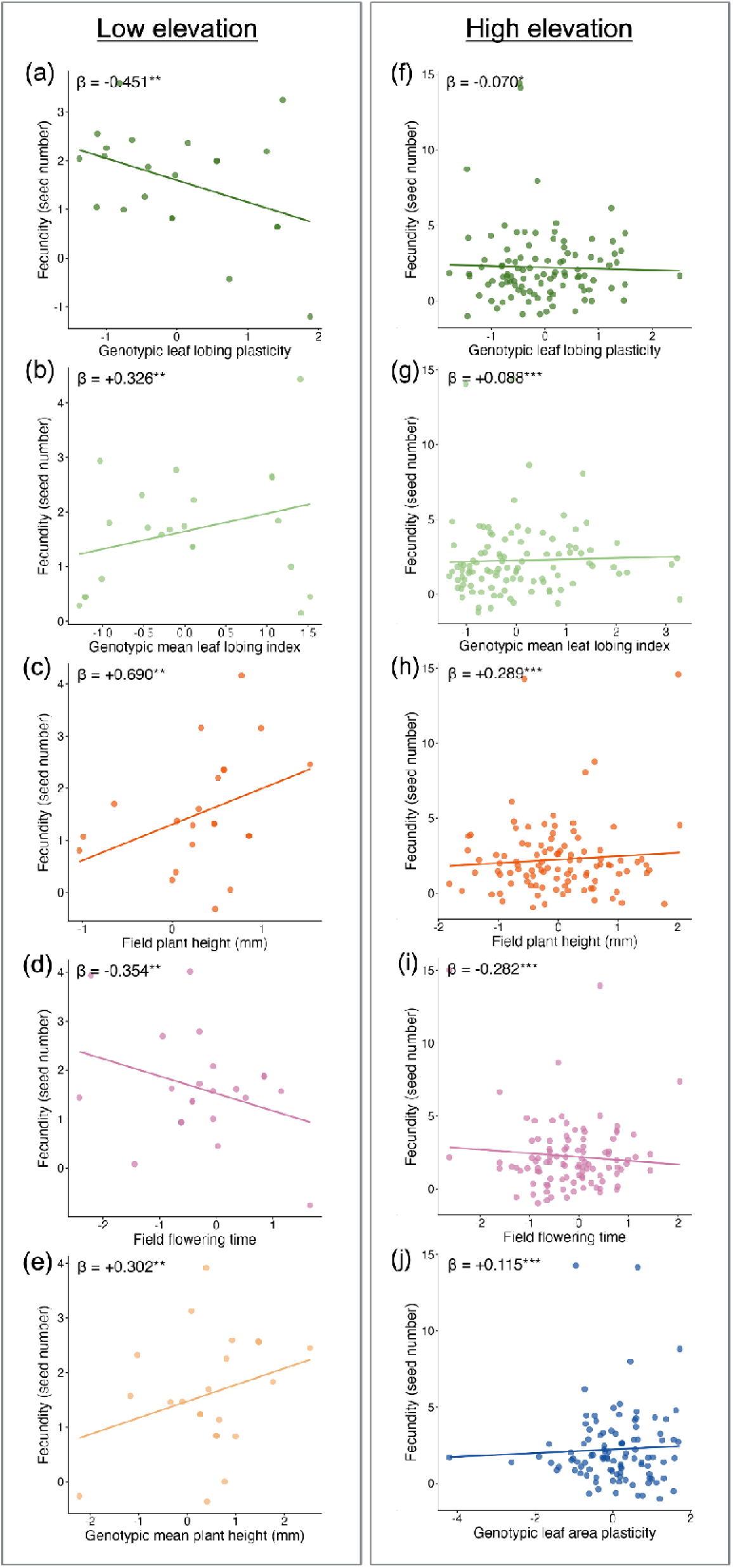
Significant directional fecundity selection gradients for traits at low and high-elevations, using partial residuals from multiple regression of zero-truncated Poisson models (Breheny & Burchett, 2017). The fitted curves show linear regressions at low and high-elevation in statistically significant traits: **a&f**) genotypic leaf lobing plasticity, **b&g**) genotypic mean leaf lobing, **c&h**) field-measured plant height, **d&i**) field-measured flowering time, **e**) genotypic mean plant height, and **j**) genotypic leaf area plasticity (Table 5). Non-significant selection gradient visualizations can be found in Fig. S3.

**Table 5.**
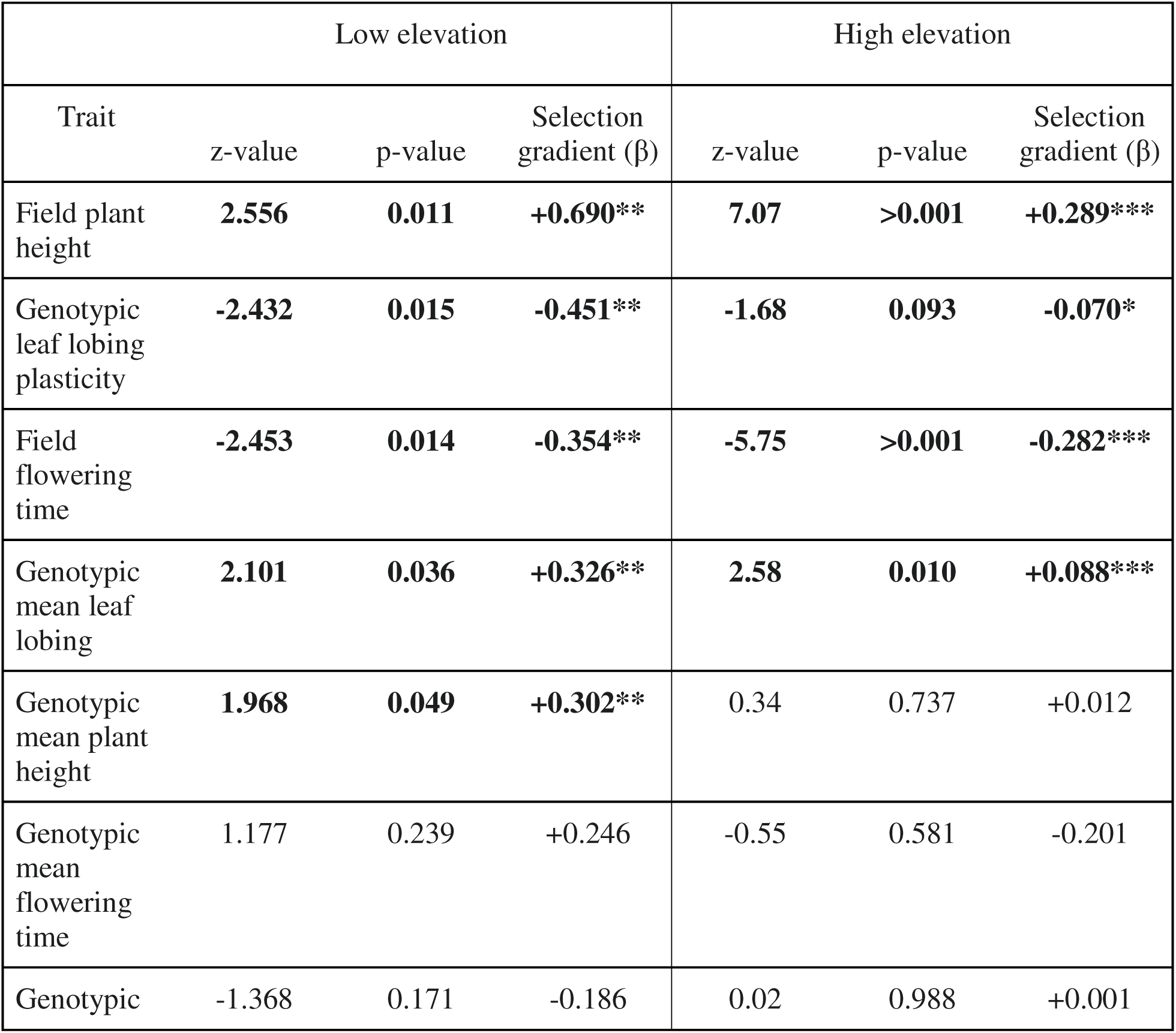
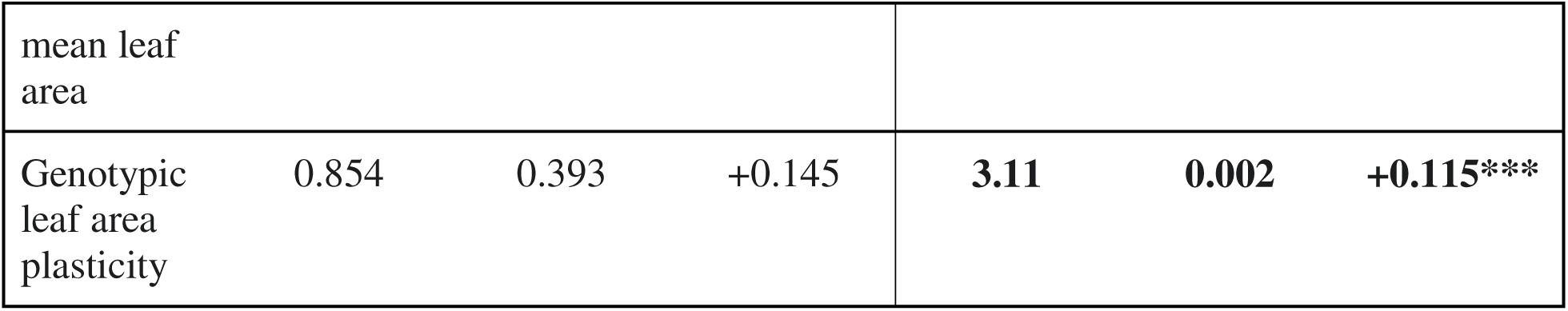
Trait influence on number of seeds produced, based on zero-truncated Poisson models run on low- and high-elevation datasets. Selection gradient (β) represents the strength and direction of selection on a trait. For low-elevation traits, numDF = 1, denDF = 18. At high-elevation, numDF = 1, denDF = 96. See Fig. 4 for visualization of these results. * p < 0.10, ** p < 0.05, *** p < 0.01

### Selection favors low-elevation leaf shape plasticity across all elevations

We ran an ANOVA on a linear model to test for effects of directional plasticity expression, experimental garden elevation, and their interaction on fecundity across elevations. Plasticity groups were defined as low-elevation (<-0.1), none (±0.1 from zero), or high-elevation (>0.1). We found that plants with low-elevation plasticity exhibited significantly higher seed output on average across gardens of different elevation than those with no plasticity (β= -2.9, t_537_ = -2.63, p = 0.009). Average fitness for high-elevation plasticity genotypes across elevations was marginally decreased compared to low-elevation plasticity genotypes (β= -1.57, t_537_ = -1.75, p = 0.081). Fitness was also significantly reduced at the low-elevation site for all genotypes (β= - 6.44, t_537_ = -5.02, p < 0.0001).

To further probe the relationship between leaf shape plasticity and fecundity and to clarify which direction of plasticity expression was under selection within each elevation, we conducted post hoc comparisons of estimated marginal means. Pairwise comparisons were conducted using the *emmeans* package in R on our zero-truncated Poisson models. We used asymptotic z-tests with degrees of freedom treated as infinite. At low-elevation, genotypes with low-elevation plasticity produced significantly more seeds than those with no plasticity (β = +1.059, z = 2.512, p = 0.032) or high-elevation plasticity (β = +0.631, z = 2.549, p = 0.029; Fig. **5a**). Genotypes with high-elevation plasticity and no plasticity did not differ in fitness (z = 1.208, p = 0.449). At high-elevation, genotypes with low-elevation plasticity similarly showed the highest fecundity compared to those with no plasticity (β = +0.756 z = 4.672, p < 0.0001) or high-elevation plasticity (β = +0.238, z = 3.350, p = 0.002; Fig. **5b**). However at high-elevation, high-elevation leaf shape plasticity also significantly increases fitness compared to no plasticity, which differs from our results at low-elevation where fitness of RILs with high-elevation plasticity is statistically equivalent to those with no plasticity (β = +0.518, z = 3.173, p = 0.004).

**Figure 5.**
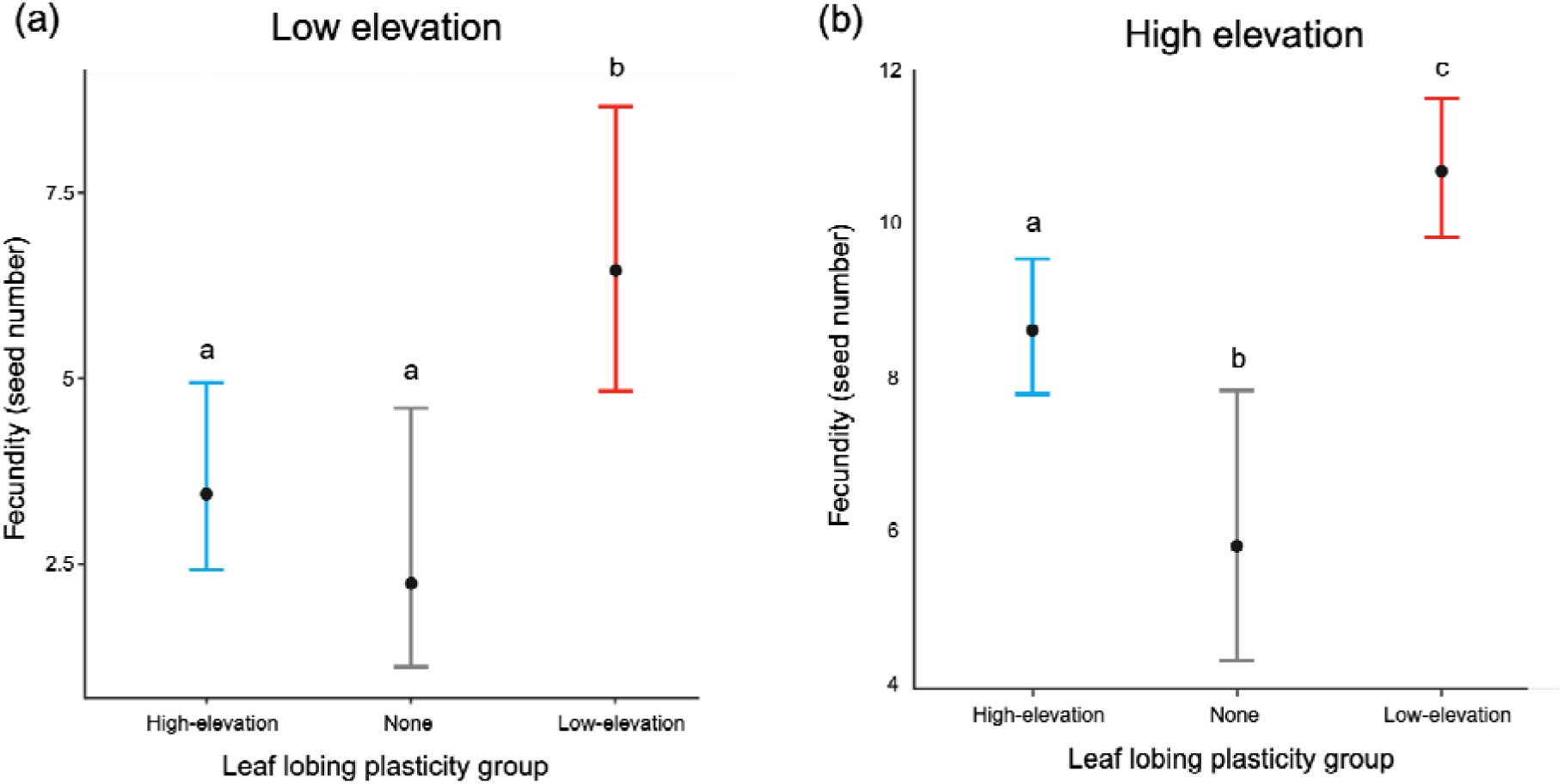
Further examination of leaf shape plasticity’s effect on plant fecundity at **a**) low-elevation and **b**) high-elevation. Asymptotic z-tests revealed significant impacts on fitness based on the direction of plasticity. Black dots indicate estimated marginal means, while error bars show a 95% confidence interval. Low-elevation leaf shape plasticity increased fecundity, and no plasticity had the lowest fecundity in both common gardens (Table 6).

**Table 6.** Influence of leaf lobing plasticity direction on fecundity, based on a post-hoc test from our zero-truncated Poisson models at low and high-elevation. We used asymptotic z-tests with degrees of freedom treated as infinite. * p < 0.10, ** p < 0.05, *** p < 0.01

|  | Low elevation |  |  | High elevation |  |  |
| --- | --- | --- | --- | --- | --- | --- |
| Comparison | z-value | p-value | Selection gradient ($\beta$) | z-value | p-value | Selection gradient ($\beta$) |
| High-elevation plasticity vs. no plasticity | 1.208 | 0.499 | +0.433 | **3.173** | **0.004** | **+0.518\*\*\*** |
| Low-elevation plasticity vs. no plasticity | **2.512** | **0.032** | **+1.059\*\*** | **4.672** | **<0.0001** | **+0.756\*\*\*** |
| Low-elevation<br>plasticity vs. High-elevation plasticity | 2.549 | 0.029 | +0.631** | 3.350 | 0.002 | +0.238*** |

## Discussion

In this study, we used a reciprocal transplant experiment between low- and high-elevation sites in the Sierra Nevada to test for selection on phenotypic plasticity and other traits that differ across elevation in *M. laciniatus*. We transplanted RILs that varied in their degree of leaf lobing plasticity, a trait previously shown to exhibit repeated, and potentially adaptive, shifts based on altitudinal clines in this species (Love & Ferris, 2024) across high and low-elevation gardens and performed genotypic selection analysis. By measuring survival and fecundity in 4,950 individuals across five common gardens, linking field-measured fitness to genotypic trait values, we found evidence of adaptive lag where low-elevation *M. laciniatus* have the highest fitness, selection for earlier flowering time and greater plant height across elevations. We also discovered positive selection on mean leaf shape consistent with adaptation to rocky outcrop environments, and that the low-elevation direction of phenotypic plasticity is adaptive across *M. laciniatus’s* altitudinal range.

### Environmental drivers of survival across the growing season

Our findings reveal a dynamic and heterogeneous survival landscape shaped by interacting abiotic stressors, and provide a mechanistic foundation for interpreting spatial variation in selection on plasticity traits in this system. Survival of experimental plants declined sharply over the course of the growing season, a pattern driven by interacting abiotic factors. The strongest predictor of survival was time since planting, underscoring the importance of temporal environmental stress, particularly the progressive drying and warming typical of montane Mediterranean ecosystems (Geber & Griffen, 2003; Heschel *et al*., 2004). Soil moisture significantly modulated the effect of time, indicating that drought was not only a main effect but interacted dynamically with seasonal progression. This aligns with findings in other montane species where early-season moisture pulses are critical to establishment, and later desiccation dramatically reduces fitness components (Franks *et al*., 2007; Angert *et al*., 2011; Ferris & Willis, 2018; Tataru *et al*., 2023).

We also found strong main and interactive effects of soil temperature, which further influenced survival, particularly in the context of soil moisture. This supports a multivariate abiotic stress environment, where temperature and moisture jointly limit survival rather than acting independently (e.g., Sherrard & Maherali, 2006; Wadgymar et al., 2018). As the season progresses, the surface temperature in hotter and drier sites may exceed physiological tolerance thresholds and accelerate desiccation, contributing to heightened mortality under compounded abiotic stress. This is particularly relevant in shallow-rooted environments typical of *M. laciniatus*.

Finally, elevation-level differences in survival reflect distinct abiotic trajectories across the elevation gradient. Our low-elevation sites had lower survival to flowering compared to high-elevation sites. Higher temperatures and lower soil moisture reduced survival overall, but their impacts varied with elevation. Low-elevation plants showed reduced benefit from soil moisture, as one low-elevation site maintained high moisture throughout the season while still facing a rapid decline in survivorship (Fig. **2a****&b**). High-elevation plants experienced greater mortality under hot conditions, coinciding with rapid drying from a mid-summer heatwave, while low-elevation plants experienced the greatest increases in temperature throughout their growing season. This variation in environmental trajectories across elevation reinforces the importance of considering local environmental context in models of adaptation and plasticity evolution (Ghalambor *et al*., 2007; Richardson *et al*., 2014). In particular, differences in how temperature and moisture interact across sites may create divergent selection regimes on traits that confer tolerance or avoidance strategies.

### Instead of local adaptation, parental genotypes show evidence of adaptive lag

Classic local adaptation theory predicts that local genotypes should exhibit higher fitness in their home environments relative to foreign genotypes (Kawecki & Ebert, 2004). We found that the low-elevation *M. laciniatus* genotype has higher fitness in both its local low-elevation habitat and at high-elevation. While fecundity between high and low parental genotypes was not significantly different, the low-elevation parent showed significantly greater survival at high-elevation, leading to increased total fitness. This differs from our predictions where we expected local vs. foreign genotypic advantage in both elevation environments. Our results suggest that the low-elevation *M. laciniatus* genotype is more robust across habitats rather than locally specialized and that the high-elevation genotype is maladapted to its current environmental conditions.

Similar patterns of asymmetric performance across environments have been observed in other plant systems and may reflect life-history trade-offs or gene flow asymmetries (Hereford, 2009; Wadgymar *et al*., 2017; Kooyers *et al*., 2019). For example, Kooyers *et al*. (2019) found that populations of *Mimulus guttatus* from California sites had higher fitness at both high and low-elevation gardens than local populations in the Cascades, OR. This indicated an adaptational lag where southern *M. guttatus* populations, historically experiencing higher temperatures and greater drought, had higher fitness in northern gardens than local genotypes under current climate scenarios. Here, the fitness of our high-elevation *M. laciniatus* may reflect adaptation to past climatic regimes, whereas the low-elevation genotype, having adapted to historically lower snowpacks and higher temperatures, may have broader physiological tolerance equating to better performance under contemporary climatic conditions. This result indicates that future upslope migration is possible and may be a likely outcome of adaptation to continued climate change in this species.

### Large plant size, early flowering, and greater leaf lobing are adaptive across *M. laciniatus*’s geographic range

Our field study found selection on a number of different traits, with many being adaptive across both low- and high-elevations (Fig. 4). Field plant height was under positive selection across both elevations; positive selection on plant height aligns with long-standing evidence that increased plant size is associated with higher fitness in annual species due to greater resource acquisition and reproductive potential (Mitchell-Olds, 1996; Stanton *et al*., 2000). Flowering time was under strong negative selection across both environments, where earlier-flowering plants had higher fecundity. This pattern is seen repeatedly in field-based annual plant studies including previous studies in *M. laciniatus* (Ferris & Willis, 2018; Tataru et al., 2023; Dong et al., 2025), where earlier flowering allows plants to reproduce before the onset of seasonal drought (Hall & Willis, 2006; Franks *et al*., 2007). These consistent patterns across altitudinal habitats suggest that despite environmental heterogeneity across *M. laciniatus*’ geographic range, some traits are under uniform directional selection. This may reflect overarching constraints of the Mediterranean climate regime and *M. laciniatus*’ unique rocky outcrop habitat, where rapid seasonal drying creates a narrow reproductive window (Fig. 2). That we detect similar selection pressures across elevation also highlights that even in highly variable environments, some functional strategies (e.g. fast, early growth and reproduction) are broadly advantageous.

Our previous common garden study (Love & Ferris, 2024) found repeated clines in leaf lobing with high-elevation populations of *M. laciniatus* having the most highly lobed leaves, leading us to hypothesize that we would find spatially varying selection on mean leaf shape across elevation in our field study. Instead, mean leaf lobing was another trait under positive selection across low- and high-elevations. Genotypic selection for greater leaf lobing was in fact stronger in the low-elevation garden than at high-elevations (Fig. **4b****&g**), opposite of our predictions based on clinal variation in natural populations. Phenotypic selection for increased leaf lobing in *M. laciniatus’s* rocky outcrop environment has been found previously (Ferris & Willis 2018) and is in line with lobed leaf shape’s hypothesized function in efficiently regulating leaf temperature and water loss in extreme environments (Nicotra *et al*., 2011). Interestingly, leaf area plasticity was only significant at high-elevation. Leaf size is also predicted to modify heat and gas exchange rates between the leaf and environment, with smaller leaves having a reduced boundary layer compared to larger leaves, allowing them to reach air temperature more efficiently through convection (Schuepp, 1993). Given our results we conclude that both leaf shape and size are likely tied to functional strategies for coping with the abiotic stressors characteristic of rocky alpine environments, with plasticity in leaf size more beneficial under high-elevation conditions. These results align with findings from *M. guttatus* as well as wild tomato (*Solanum pennellii*) where plasticity in leaf morphology has been associated with higher fitness in fluctuating environments (Sultan, 2000; Chitwood *et al*., 2013; Friedman *et al*., 2015).

### Genotypic selection for plasticity varies across elevation

The central question of this study was whether plasticity in leaf lobing is locally adaptive across environments, as suggested by our previous work where we found repeated altitudinal clines in plasticity, with low-elevation populations having less leaf shape plasticity than high-elevation populations (Love & Ferris, 2024). Our present findings indicate that the fitness consequences of plasticity in leaf lobing are broadly beneficial, while still being environmentally dependent. In our binomial model, testing for the effect of a trait on the probability of survival to seed production, we found genotypic selection for high-elevation leaf shape plasticity in our high-elevation gardens and no selection on plasticity in low-elevation gardens (Table 4). This pattern is consistent with our prediction that plasticity aligned with the local elevation environment confers a reproductive advantage. Similar environment-specific selection on plasticity has been documented in other systems. For example, *Impatiens capensis* populations exhibited density- and site-dependent selection on plastic shade-avoidance traits, where plastic individuals performed better only under local light environments (Donohue *et al*., 2000). Likewise, in *Brassica rapa*, plastic genotypes expressed fitness costs in high-density environments, demonstrating that selection on plasticity can depend on competitive context (Dechaine *et al*., 2007). These studies illustrate that plasticity is often beneficial only under certain environmental pressures, rather than universally adaptive.

In fecundity selection, we found RILs with low-elevation plasticity (negative values) were selected for at both low and high-elevations (Table 5, Fig. 4**a****&f**). A post-hoc selection analysis of plasticity revealed that this pattern was driven primarily by the strong fitness of genotypes exhibiting low-elevation plasticity. However, in high-elevation gardens both low- and high-elevation plasticity RIL genotypes had greater fitness than non-plastic genotypes (Table 6, Fig. 5). To measure genotypic values of mean leaf shape and leaf shape plasticity we used genotype means derived from controlled growth chamber and greenhouse environments which follows from genotypic selection theory and allows us to estimate selection on trait values that reflect heritable, environment-dependent expression (Rausher, 1992; Stinchcombe & Rausher, 2001). One possible limitation of this method is that trait expression was not quantified in the same environments where fitness was measured. However, our growth chamber treatments quantifying plasticity were designed to simulate photoperiod conditions at the extremes of the species’ elevational range, and prior work has shown that these treatments successfully elicit biologically meaningful variation in leaf shape (Love & Ferris, 2024). Our use of replicated RILs across field sites strengthens the inference of selection on genotype-level traits and plasticity, even if not all traits could be measured directly in the field.

Comparisons to other examples of selection in the literature may help interpret the unexpected pattern of low-elevation plasticity genotypes having the highest fecundity across elevations. In *Arabidopsis thaliana*, Wilczek et al. (2009) conducted a broad European reciprocal transplant experiment and found that phenotypic plasticity in life history traits (especially flowering time) varied among genotypes and environments but did not always translate into higher fitness in home environments, highlighting the intricacy of fitness plasticity relationships. Similarly, work on poplar (*Populus spp.*) shows that variation in plastic responses of leaf and stem traits can influence growth and competitive success under different environmental conditions, but that plasticity can be maintained by fluctuating selection rather than directional local adaptation (Cope *et al*., 2021). In this context, our observation that low elevation plasticity genotypes perform well everywhere may reflect a similar scenario where plastic responses that are more generalist in nature contribute to fitness across a range of conditions.

The broader literature suggests that plasticity is most likely to be maintained when environments fluctuate spatially and temporally, such that no single trait value is optimal everywhere (Crispo, 2008; Snell-Rood & Ehlman, 2021). In high-elevation environments with unpredictable timing of snow melt and snow fall, balancing selection may maintain both directions of leaf lobing plasticity based on conflicting selection gradients from interannually varying conditions (Dong *et al*., 2025). It is possible that temporally variable selection, known to be a persistent force in *M. laciniatus*’s rocky outcrops due to dramatic interannual climatic fluctuations (Tataru *et al*., 2023; Dong *et al*., 2025), maintains genetic variation in plasticity direction. Our experiment, performed during an average snowpack year but where snowmelt happened rapidly, may have caught conditions that give low-elevation plasticity genotypes a universal advantage. It is also possible that, like our results in the relative fitness of high and low *M. laciniatus* parental genotypes, this result indicates a mismatch between historical and current selection pressures creating adaptive lag in phenotypic plasticity as well as parental fitness (Kooyers *et al*., 2019). Perhaps as the climate continues to warm and the timing of snowmelt moves earlier, creating exposure to shorter day lengths during the growing season at high-elevations, we will see an increase in frequency of low-elevation plasticity in high-elevation *M. laciniatus* populations. Previously we found that high-elevation populations showed strong high-elevation leaf lobing plasticity, but our fecundity selection analysis does not provide evidence for this pattern being locally adaptive. However, genetic variation within natural high-elevation populations still exists such that some high-elevation genotypes exhibited the low-elevation direction of plasticity in a previous experiment (Love & Ferris 2024). Low-elevation environments may favor low-elevation plasticity, where more lobed leaves are produced under local shorter photoperiods, as a developmentally labile means to produce lobed leaves uniquely found in rocky outcrop *Mimulus* species (Ferris *et al*., 2015) and are under positive selection in *M. laciniatus’s* unique habitat (Ferris & Willis, 2018).

### Minimal costs of phenotypic plasticity in a native field environment

Although plasticity is often assumed to be adaptive, many theoretical studies have asked whether plasticity itself is costly (Via *et al*., 1995; DeWitt, 1998). These costs may occur via the energetic expense of developing or maintaining a plastic response, or through maladaptive mismatches under certain conditions (Auld *et al*., 2010). In our field experiment, we found that leaf shape plasticity was broadly beneficial, enhancing fitness across both elevation environments. However, the adaptive value of plasticity depended on the direction of the trait expression: genotypes expressing low-elevation plasticity (i.e., negative reaction norms) had the highest fecundity at both low and high-elevation, while genotypes with high-elevation plasticity (i.e., positive reaction norms) only had higher viability at high-elevation. This pattern suggests asymmetric selection on plasticity and raises the possibility of context-dependent costs, as we detected differential selection on three different directions of reaction norm slope (negative, neutral, and positive) in one plastic trait.

Yet, it is important to note that we did not observe plastic genotypes performing worse than non-plastic genotypes in any environment, and thus did not detect strong evidence of costs of plasticity. This aligns with a large body of literature showing that costs of plasticity are difficult to detect in field conditions, only emerging under specific or stressful contexts (Van Kleunen & Fischer, 2005; Callahan *et al*., 2008; Matesanz *et al*., 2010). For example, in *Boechera stricta*, Wagner et al. (2018) found that plasticity in defense chemistry was associated with greater fitness across contrasting habitats but did not show consistent selection within any single environment, reinforcing the idea that plastic responses can be beneficial across multiple contexts without local tuning. Empirical tests across diverse taxa often fail to find consistent fitness costs associated with plastic responses, especially when plasticity leads to environmentally appropriate phenotypes (Ghalambor *et al*., 2007; De Lisle & Rowe, 2023). Our results contribute to this literature by demonstrating that, in a median-snowfall year with a perhaps more relaxed selection regime than drought years, leaf shape plasticity conferred fitness benefits rather than repercussions. Adding to this finding, we saw that the direction of plasticity expression mattered, emphasizing that adaptive plasticity is not always symmetric across different environments. We thus conclude that in *M. laciniatus*, costs likely manifest not from the mechanistic presence of plasticity itself, but from environmental mismatches in plastic responses.

## Conclusion

Taken together, our findings offer new insight into how natural selection acts on phenotypic plasticity in the wild. We contribute empirical evidence from a large-scale reciprocal transplant experiment, showing an adaptive lag of high-elevation genotypes, that plastic genotypes are always more fit than non-plastic genotypes, and that local directions of plasticity are sometimes, but not always, more beneficial. Therefore, trait-environment mismatches may occur even in a species where plasticity is common. The adaptive significance of phenotypic plasticity has long been debated in evolutionary ecology, in recent years becoming especially prominent in light of the role plasticity could play in the race against ongoing global climate change. We find that plasticity in one trait may be both adaptive and maladaptive, depending on the environmental context. As high-elevation plant populations continue to experience historically earlier springs, growing under warmer temperatures and shorter photoperiods, we may see an upslope migration of leaf shape plasticity genotypes in *M. laciniatus*. Our work advances understanding of the evolutionary dynamics of plasticity in heterogeneous landscapes and highlights the value of integrative approaches to link trait variation, phenotypic plasticity, and fitness in the wild.

## Acknowledgements

We thank lab technician Juj Sullivan and undergraduate student Lillian Milgram for their extensive help planting common gardens in Yosemite. We also thank Chloe Love, Catherine Ogoma, Gabrielle Watson, Diana Tataru, Caroline Dong, and Whitney Murchison-Kastner for additional fieldwork assistance, support, and advice. We thank the Sexton Lab at UC Merced and the UC Merced greenhouse staff for assisting us with the use of their facilities. We also thank Breezy Jackson and Marlon Spinneberg of the Sierra Nevada Research Station - Yosemite Field Station (doi:10.21973/N3V36C) for housing coordination and assistance. Thank you to the National Park Service for research support and permitting (permit #YOSE-2024-SCI-005). Thank you to undergraduate students Jessie Lampert, Sydney Sarkisian, and Madi Kelso for help with greenhouse and lab data collection. Finally, thank you to Emily Josephs, Sunshine Van Bael, Alex Gunderson, and the Ferris Lab for their thoughtful comments on the manuscript.

This work was funded by an American Philosophical Society Lewis and Clark Grant for Exploration and Field Research, a Tulane University EEB Graduate Research Grant, and a Tulane University Connolly Alexander Institute for Data Science Graduate Research Grant awarded to JML. This manuscript was made possible in part by a residency by JML at A Studio in the Woods (Tulane University). This research was supported by the National Institute of General Medical Sciences of the National Institute of Health (NIH) under Award Number R35GM138224 awarded to KGF. The content is solely the responsibility of the authors and does not necessarily represent the official views of the NIH.

We acknowledge the long-standing stewards of the unceded land upon which our field work was conducted: the Southern Sierra Miwuk Nation, Bishop Paiute Tribe, Bridgeport Indian Colony, Mono Lake Kutzadikaa, North Fork Rancheria of Mono Indians of California, Picayune Rancheria of the Chukchansi Indians and the Tuolumne Band of Me-Wuk Indians. As scientists, we strive to take responsibility for the impacts of colonialism in our field and move forward with respect and support of indigenous movements and knowledge.

## Author contributions

JML and KGF conceptualized the research, funded the research, and wrote the manuscript. JML performed all experiments, fieldwork, data collection, and data analysis. KGF provided supervision and other resources.

## Data availability

Data and code are available on GitHub at the following repository: https://github.com/jill-love/Love-Ferris-2026

