## supplemental for "Phenotypic plasticity is broadly adaptive across an elevation gradient in the Cutleaf Monkeyflower"

Supplemental Resources for the manuscript “Phenotypic plasticity is broadly adaptive across an elevation gradient in the Cutleaf Monkeyflower”

Authors: Jill M. Love and Kathleen G. Ferris

**Supplemental Figures**

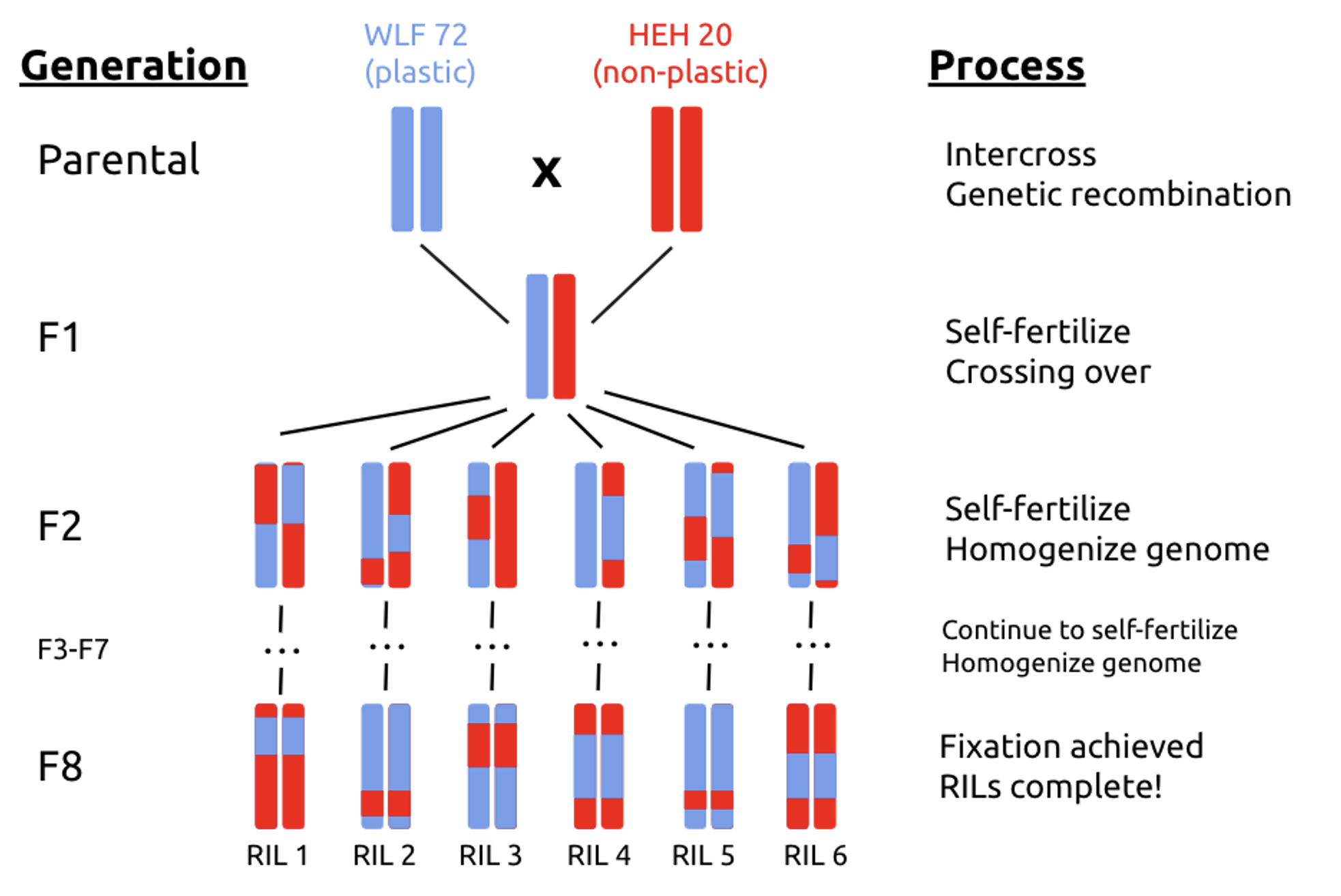

Figure S1. Crossing design of RIL population.

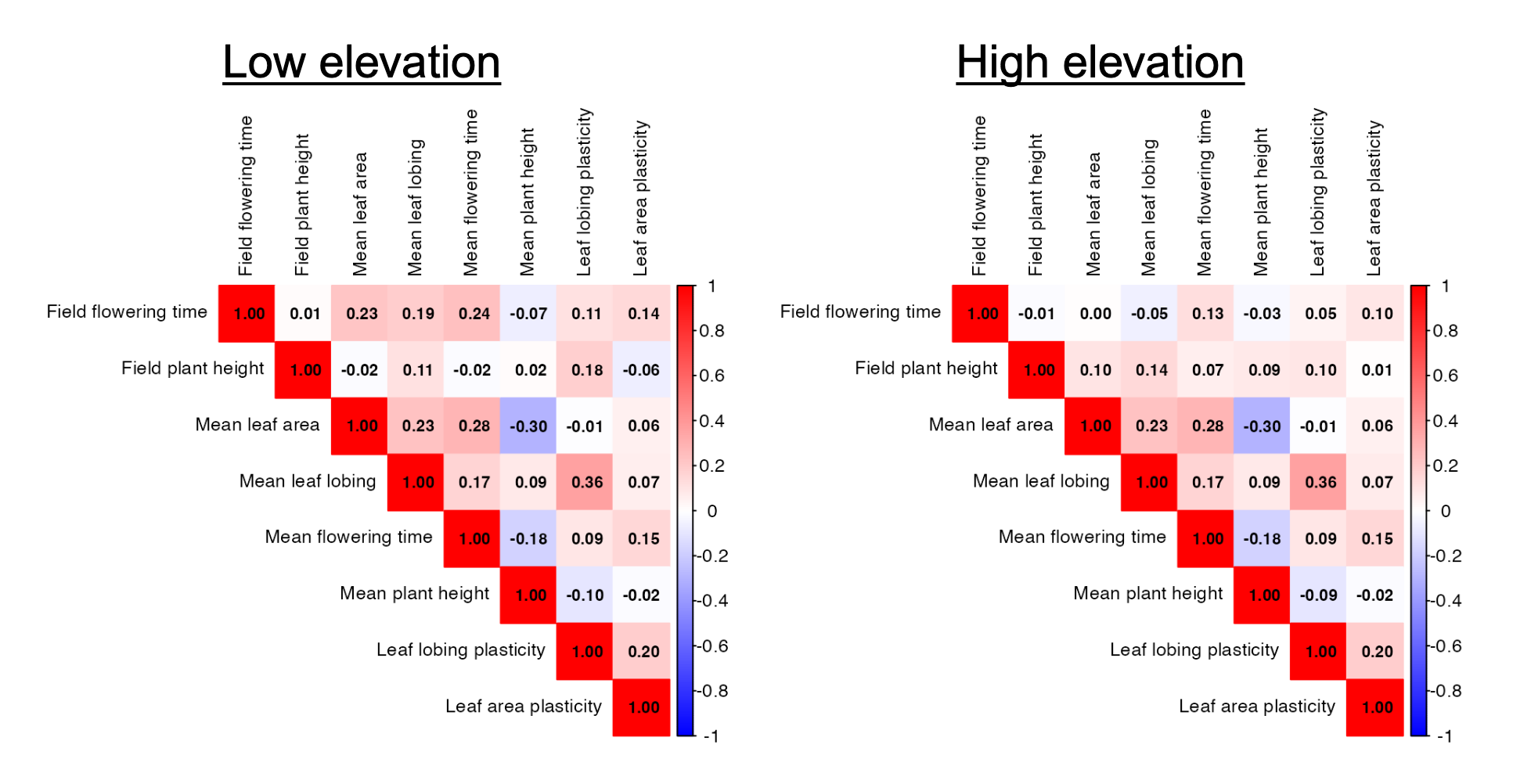

Figure S2. Correlation matrices of all traits included in genotypic selection models for low and high-elevation.

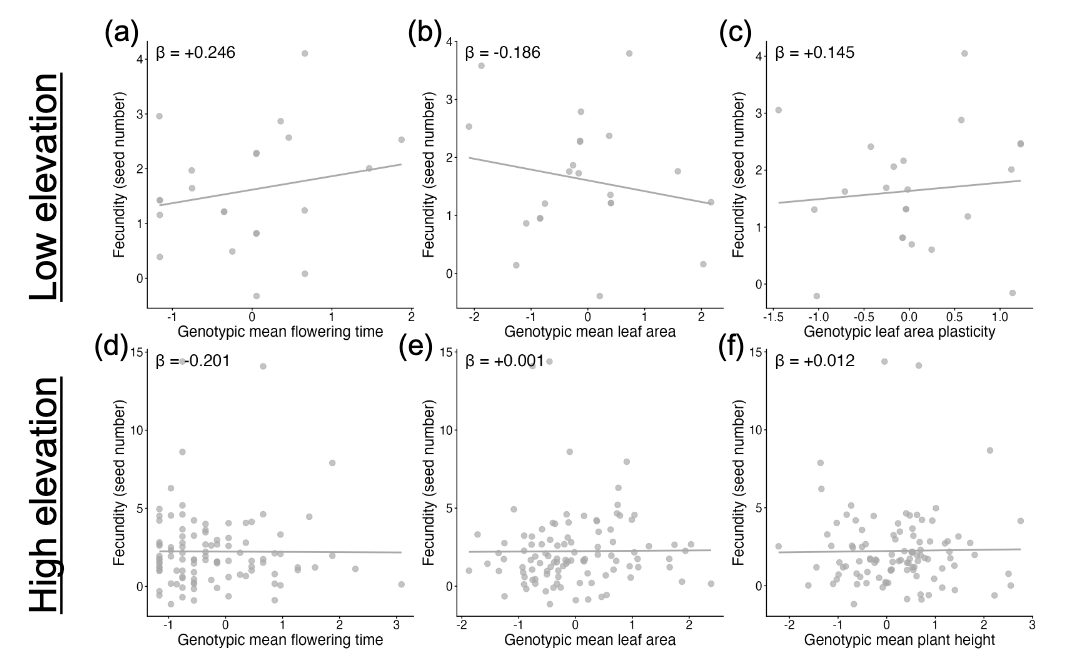

Figure S3. Non-significant directional fecundity selection gradients for remaining traits at low and high-elevations, using partial residuals from multiple regression of zero-truncated Poisson models (Breheny & Burchett, 2017). The fitted curves show linear regressions at low and high-elevation in statistically significant traits: **a&d)** genotypic mean flowering time, **b&e**) genotypic mean leaf area, **c**) genotypic leaf area plasticity, and **f**) genotypic mean plant height (Table 5).

**Supplemental Tables**

Table S1. Comparison of best-fit binomial and zero-truncated Poisson models for low- and high-elevation sites after model selection using MuMIn. Log(L) and AIC are provided to assess relative model fit.

| Model type | Elevation | Model description | Log(L) | AIC |
| --- | --- | --- | --- | --- |
| Binomial | Low | any seeds ~ field flowering time + field plant height | -33.5 | 73.0 |
|  | High | any seeds ~ genotypic mean flowering time + field flowering time + field plant height | -38.2 | 86.5 |
| Zero-truncated Poisson | Low | fecundity ~ field plant height + genotypic mean plant height + genotypic leaf lobing plasticity | -50.5 | 109.0 |
|  | High | fecundity ~ field plant height + genotypic mean plant height + genotypic leaf lobing plasticity + field flowering time + genotypic leaf area plasticity + genotypic mean leaf lobing | -445.5 | 902.9 |

Table S2. Trait influence on survival to seed production (best-fit models). For low-elevation traits, numDF = 1, denDF = 171. At high-elevation, numDF = 1, denDF = 169. Variables with n.s. were not included in the best-fit model and therefore do not have a selection gradient to report. * p < 0.10, ** p < 0.05, *** p < 0.01

|  | Low-elevation | | | High-elevation | | |
| --- | --- | --- | --- | --- | --- | --- |
| Trait | z-value | p-value | Selection gradient (β) | z-value | p-value | Selection gradient (β) |
| Field plant height | 1.628 | 0.104 | +0.578 | **3.050** | **0.002** | **+1.689***** |
| Field flowering time | -1.457 | 0.145 | -0.489 | **-2.189** | **0.029** | **-0.736**** |
| Genotypic leaf lobing plasticity | n.s. | n.s | n.s. | **1.76** | **0.078** | **+0.675*** |
| Genotypic mean flowering time | n.s. | n.s | n.s. | -1.553 | 0.121 | -0.500 |

Table S3. Trait influence on number of seeds produced (best-fit models). For low-elevation traits, numDF = 1, denDF = 18. At high-elevation, numDF = 1, denDF = 96. See Fig. 4 for visualization of these results. Variables with n.s. were not included in the best-fit model and therefore do not have a selection gradient to report. * p < 0.10, ** p < 0.05, *** p < 0.01

|  | Low-elevation | | | High-elevation | | |
| --- | --- | --- | --- | --- | --- | --- |
| Trait | z-value | p-value | Selection gradient (β) | z-value | p-value | Selection gradient (β) |
| Field plant height | **3.008** | **0.003** | **+0.638***** | **7.28** | **>0.001** | **+0.288***** |
| Genotypic mean plant height | **2.812** | **0.005** | **+0.359***** | n.s. | n.s. | n.s. |
| Genotypic leaf lobing plasticity | **-2.145** | **0.032** | **-0.263**** | **-1.84** | **0.065** | **-0.075*** |
| Field flowering time | n.s. | n.s. | n.s. | **-5.86** | **>0.001** | **-0.283***** |
| Genotypic leaf area plasticity | n.s. | n.s. | n.s. | **3.16** | **0.002** | **+0.116***** |
| Genotypic mean leaf lobing | n.s. | n.s. | n.s. | **2.84** | **0.004** | **+0.088***** |
